# Multi-pronged analysis of pediatric low-grade glioma reveals a unique tumor microenvironment associated with BRAF alterations

**DOI:** 10.1101/2024.04.05.588294

**Authors:** Shadi Zahedi, Kent Riemondy, Andrea M. Griesinger, Andrew M. Donson, Rui Fu, Michele Crespo, John DeSisto, Madeline M. Groat, Emil Bratbak, Adam Green, Todd C. Hankinson, Michael Handler, Rajeev Vibhakar, Nicholas Willard, Nicholas K. Foreman, Jean Mulcahy Levy

## Abstract

Pediatric low-grade gliomas (pLGG) comprise 35% of all brain tumors. Despite favorable survival, patients experience significant morbidity from disease and treatments. A deeper understanding of pLGG biology is essential to identify novel, more effective, and less toxic therapies. We utilized single cell RNA sequencing (scRNA-seq), spatial transcriptomics, and cytokine analyses to characterize and understand tumor and immune cell heterogeneity across pLGG. scRNA-seq revealed tumor and immune cells within the tumor microenvironment (TME). Tumor cell subsets revealed a developmental hierarchy with progenitor and mature cell populations. Immune cells included myeloid and lymphocytic cells. There was a significant difference between the prevalence of two major myeloid subclusters between pilocytic astrocytoma (PA) and ganglioglioma (GG). Bulk and single-cell cytokine analyses evaluated the immune cell signaling cascade with distinct immune phenotypes among tumor samples. *KIAA1549-BRAF* tumors appeared more immunogenic, secreting higher levels of immune cell activators and chemokines, compared to *BRAF V600E* tumors. Spatial transcriptomics revealed the differential gene expression of these chemokines and their location within the TME. A multi-pronged analysis of pLGG demonstrated the complexity of the pLGG TME and differences between genetic drivers that may influence their response to immunotherapy. Further investigation of immune cell infiltration and tumor-immune interactions is warranted.

**Key points:**

- There is a developmental hierarchy in neoplastic population comprising of both progenitor-like and mature cell types in both PA and GG.
- A more immunogenic, immune activating myeloid population is present in PA compared to GG.
- Functional analysis and spatial transcriptomics show higher levels of immune mobilizing chemokines in *KIAA1549-BRAF* fusion PA tumor samples compared to *BRAF* V600E GG samples.

**Importance of the Study:** While scRNA seq provides information on cellular heterogeneity within the tumor microenvironment (TME), it does not provide a complete picture of how these cells are interacting or where they are located. To expand on this, we used a three-pronged approach to better understand the biology of pediatric low-grade glioma (pLGG). By analyzing scRNA-seq, secreted cytokines and spatial orientation of cells within the TME, we strove to gain a more complete picture of the complex interplay between tumor and immune cells within pLGG. Our data revealed a complex heterogeneity in tumor and immune populations and identified an interesting difference in the immune phenotype among different subtypes.

## Introduction

Pediatric low-grade gliomas (pLGG) are the most common type of pediatric brain tumors accounting for approximately one third of all central nervous system (CNS) tumors (1). Surgical resection remains the first line of therapy and can be curative in some cases. For the remainder of patients, chemotherapy and sometimes radiation are needed. Although the five-year event free survival is encouraging in historical studies, 40% of pLGG patients relapse within five years of diagnosis and require additional therapy (2). Additionally, depending on the location of the tumor and the mutations they carry, pLGG can lead to significant neurologic damage, multiple recurring rounds of therapy, and in some patients, ultimately death (3). The significant morbidity and even mortality associated with pLGG highlights an urgent need for new treatment strategies.

Harnessing the immune system to treat human cancer has transformed the field of oncology but has yet to significantly impact the pLGG population. Immunotherapies have shown promising results in several tumor types. However, efficacies of such treatments vary across tumors and patients (4, 5). CNS tumors have been particularly difficult to target. Recent data suggest the CNS is an immunologically ‘distinct’ location (6). Despite this new understanding of how the immune system can interact with the CNS, studies have shown that brain tumors are able to develop immune evasion in a variety of unique ways such as sequestering T-cells in the bone marrow to encourage antigenic ignorance (7). Traditionally, CNS tumors were often found to have low numbers of tumor-infiltrating lymphocytes and other immune effector cells (8). This is felt to be primarily related to the tumor/immune microenvironment in the CNS which has evolved to tightly regulate inflammation that could have detrimental effects in the enclosed cranial space (9). Studies investigating immune infiltrating cells in adult glial tumors have uncovered tumor suppressive factors including myeloid-derived suppressor and T-regulatory cells, as well as immune suppressive metabolites. As a result, researchers have developed strategies to target inhibitory cells and molecules and utilize the power of the immune system to treat adult CNS tumors (10). In previous studies from our group, we have demonstrated different immune-phenotypes across pediatric brain tumor pathologies with a suggestion that PA’s have a less suppressive immune environment (11). Overall, studies on the nature of tumor infiltrating immune cells in pediatric pLGG are limited but may help with improving immunotherapy outcomes in these patients.

pLGG is thought of as a single pathway disease due to the uniform up-regulation of the mitogen-activated protein kinase (MAPK) pathway (12). This is a result of mutations such as *BRAF V600E*, *NF1* and *KIAA1549-BRAF* fusion. *KIAA1549-BRAF* fusion is the most frequent molecular alteration, identified in 30-40% of pLGG (12). This mutation is significantly enriched in PA and in tumors arising in the posterior fossa. PA have highly circumscribed histology and normally arise in surgically amenable locations. However, when these tumors arise in deeply seated areas of the brain where complete surgical resection is not possible, progression becomes more common (12).

The prevalence of *BRAF V600E* tumors varies notably depending on the histology and the location of the tumor. pLGG that frequently harbor *BRAF* V600E include pleomorphic xanthoastrocytoma (40-80%), diffuse astrocytoma (30-40%) and ganglioglioma (25-45%) (12). Overall, *BRAF V600E* tumors have been associated with the lowliest overall and progression free survival compared to other pLGG. Especially in the context of co-occurring *CDKN2A* deletions, *BRAF V600E* pLGGs are significantly more likely to transform into higher-grade tumors, often decades following the initial diagnosis (13).

Whole genome and RNA sequencing (RNA-seq) technologies have been used to study gene expression patterns in many tumors, including pLGG. These studies identified additional mutations in *FGFR1*, *PTPN11*, and *NTRK2* fusion genes (14). In addition, the combination of RNA-seq and copy number variation data has identified novel fusion partners with the *BRAF* gene (15). These data have brought a greater understanding of pLGG biology and advances to therapy including the use of MAPK pathway inhibitors. Although these inhibitors have decreased morbidity and improved survival, there is already evidence for developing resistance (16). Additionally, these studies were performed on snap frozen, bulk tumor samples which lack the resolution to capture the complexity of the TME at the single cell level.

Single cell RNA-seq (scRNA-seq) provides the opportunity to investigate gene expression profiles at the single cell level and can provide insight into the diversity of tumor infiltrating immune cells. One scRNA-seq study of six PA A2B5+ glial progenitor patient samples found these cells had a differentiated, astrocyte-like phenotype and a smaller number of cells with a proliferative, progenitor-like phenotype (17). This study noted that 40% of identified cells were immune related but there was insufficient expression of immune checkpoint genes suggesting additional study of PA associated immune cells was needed. Most recently a single-nucleus RNA seq study evaluated five ganglioglioma patient samples and found CD34+ neuroectoderm-like tumor precursor cells in addition to immune cells with myeloid and lymphoid lineage cells with potentially immune suppressive components (18). These papers offer an early glimpse into the complex nature of pLGG, but there remains a gap in understanding these tumors, the role of immune cells within them, and how these tumors compare to each other with neoplastic and non-neoplastic components.

To attempt a broader understanding of pLGG and the connected tumor microenvironment (TME) we investigated a panel of 23 patient samples using a multi-pronged approach including scRNA seq, single cell and bulk cytokine analyses and spatial transcriptomics (ST). Together, these data provide a clearer picture of a complex tumor heterogeneity between pLGG subtypes.

## Results

### Patient sample characteristics

Following informed consent, samples were obtained from pediatric low-grade glioma patients who underwent tumor resection during 2011-2019 (Table 1). Samples were disaggregated at the time of surgery and viably banked at Morgan Adams Pediatric Neuro-Oncology Program Biobank. All samples were from initial diagnosis except one of the GG samples (UPN# 1492) which was from the first recurrence after therapy (the patient was treated with dabrafenib for two years). The median age at diagnosis was 10 years (range, 1-17). The most common diagnosis was PA (n=14) followed by GG (n=7) and not otherwise specified undefined low-grade gliomas (n=2). Tumor locations varied across samples, although the most common site was posterior fossa. The majority of PA patients carried *KIAA1544:BRAF* fusion alteration (n=13) and one PA had a *FGFR1* mutation. Six of seven GG patients carried *BRAF* V600E mutation and 1 patient carried a *FGFR3:TACC3* fusion mutation. One of the undefined LGG patients had no identified mutations and the other carried a *FGFR3:TACC3* fusion. We confirmed the mutation status of all tumors by genetic profiling of bulk tissues.

**Table 1:**
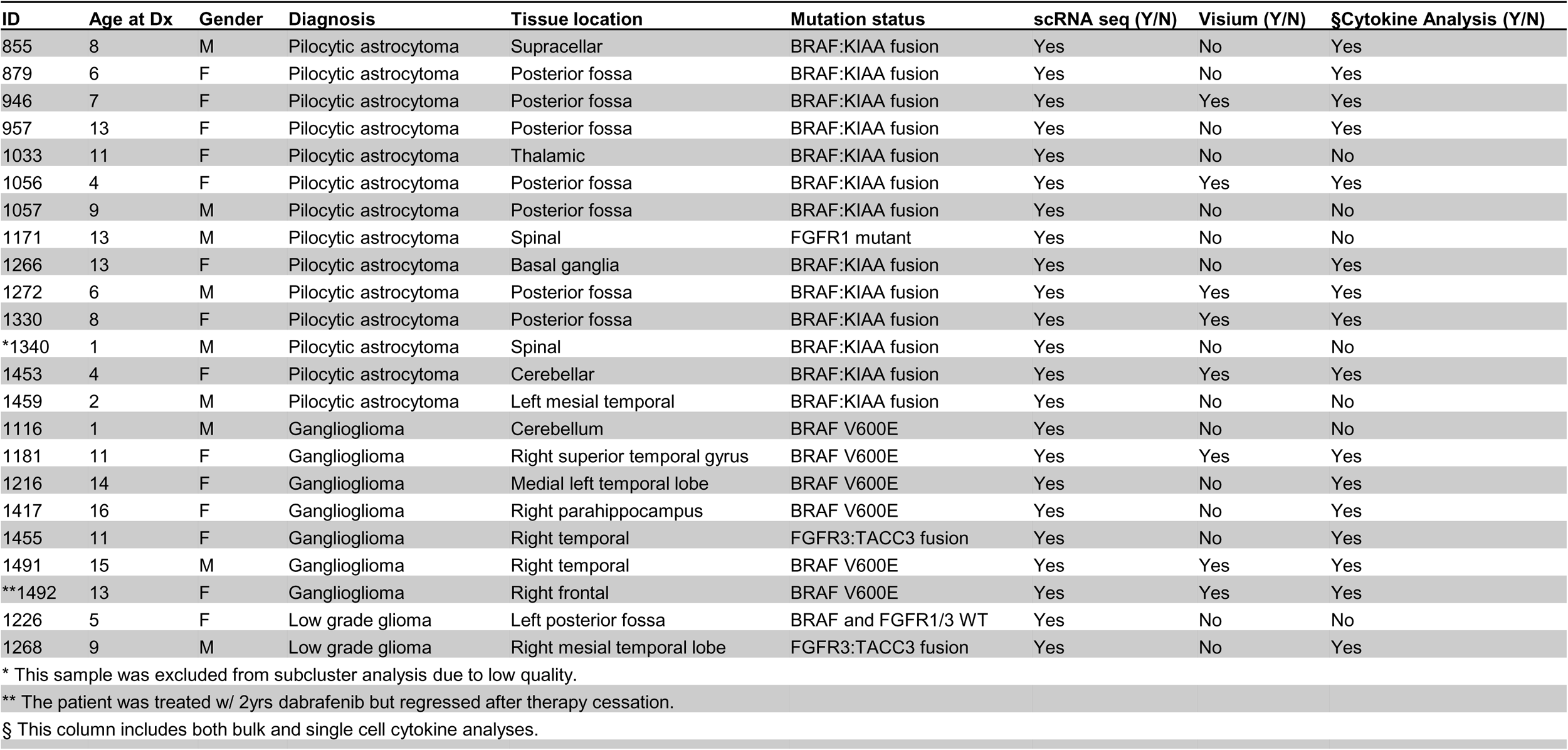
Summary of pLGG sample characteristics.

### scRNA-seq of pediatric low-grade glioma (pLGG) demonstrates neoplastic and non-neoplastic subgroups

To determine the extent of cellular heterogeneity and complexity of pLGG and its microenvironment, we performed scRNA-seq on 23 pLGG primary patient samples. Cell Ranger (10X Genomics) and Seurat Analyses were used to filter and normalize cells, resulting in 26,029 cells that passed quality controls (Fig 1A). These cells were projected as UMAP plots, revealing multiple clusters shared across different samples (Figure 1A-B). Harmony alignment was used to correct for inter-sample variations from experimental or sequencing batch effects. Harmony alignment revealed multiple clusters harboring gene expression profiles of both neoplastic and non-neoplastic clusters. Non-neoplastic clusters comprised both myeloid and lymphoid lineage cells (Figure 1A). Known neoplastic and non-neoplastic markers were expressed in separate clusters as expected (Supplementary Figure 1). While there were differences in the proportion of neoplastic and non-neoplastic cells in each patient sample, every individual sample included each cell type (Figure 1B). To examine the proportion of neoplastic and non-neoplastic clusters identified by scRNA seq, we performed deconvolution analyses on pLGG bulk tissue transcriptome profiles using CYBERSORTx (Supplementary Figure 2). The data from bulk tissue demonstrated a similar finding of a mix of immune and tumor cells for each sample, although the proportion of glioma cells was higher in the deconvolution analysis.

**Figure 1:**
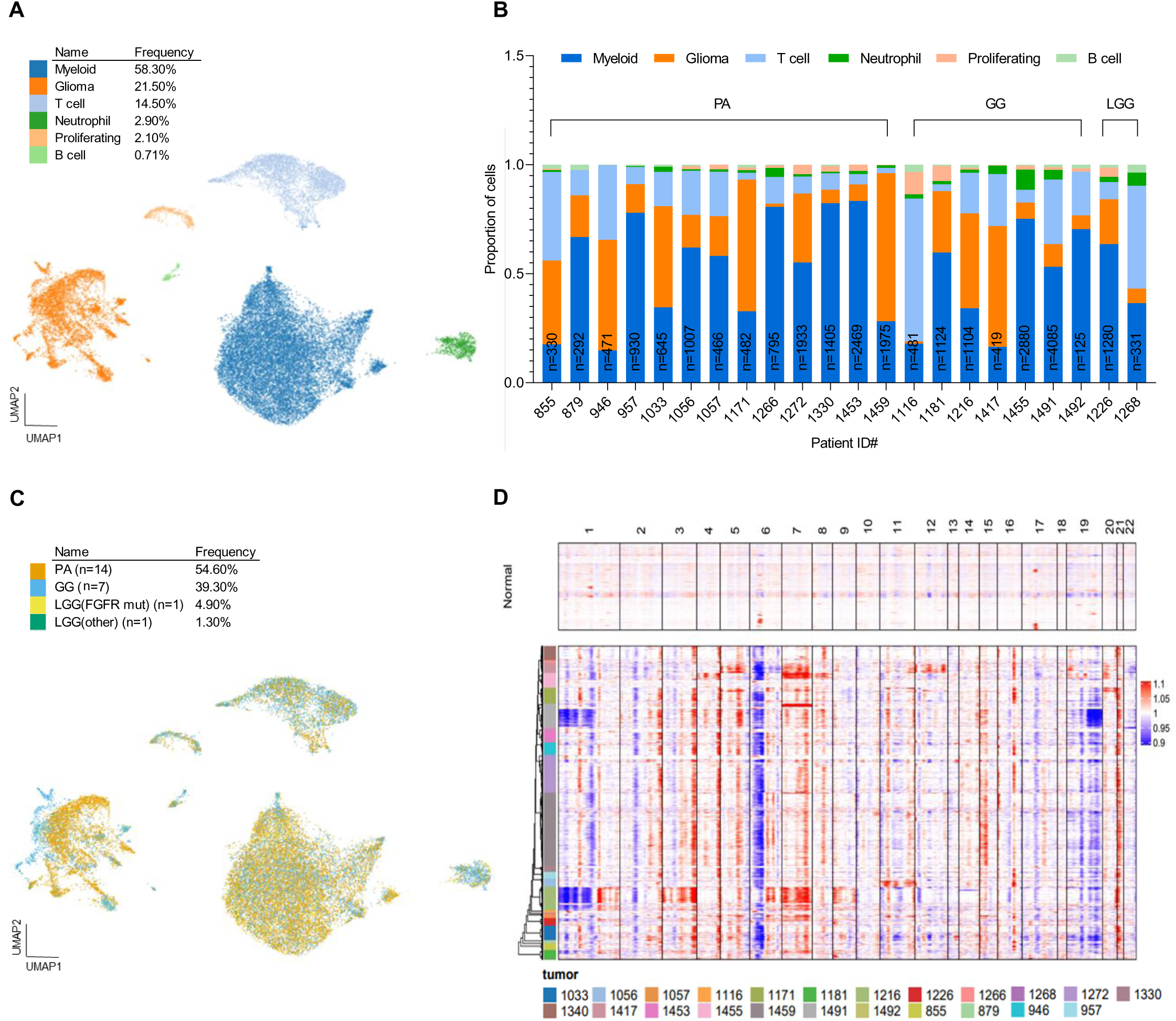
scRNA-seq analysis of pediatric low-grade glioma reveals neoplastic and non-neoplastic clusters. **A)** Harmony aligned UMAP projection of single-cell expression data of 23 pLGG samples colored by neoplastic and several non-neoplastic (myeloid, T cell, B cell, Nt, proliferating) clusters. Cluster type abundance is indicated by %frequency in the legend. **B)** Stacked bar charts show the contribution of each patient sample single cells to the neoplastic and non-neoplastic clusters. Cell counts for each sample are normalized to 1 and are color-coded according to the specific cluster. **C)** Harmony aligned UMAP projection of neoplastic and non-neoplastic clusters colored by tumor type. The number of samples and cellular abundance contributing to each tumor type is indicated in the legend. **D)** Inference of CNV (inferCNV) profiles of neoplastic and non-neoplastic PLGG single cells. Abbreviations: Nt, neutrophils; CNV, copy number variants; PA, pilocytic astrocytoma; GG, ganglioglioma; LGG, low-grade glioma; mut, mutation.

Additionally, single cells were labeled by tumor type to assess the distribution of cells that comprise different pLGG subtypes across major subclusters (Figure 1C). Non-neoplastic cells from both PA and GG showed similar distribution, although more GG cells were observed in T cell and Neutrophil clusters compared to PA cells. There were distinct differences in PA and GG tumor cell clusters (Figure 1C) suggesting a potential differential gene expression pattern.

We used copy number variations (CNVs) to distinguish pLGG cells from other cells. The most notable CNV events included losses on chromosomes 1, 6, and 19 as well as gains on chromosomes 3, 5, 7, and 21 (Figure 1D). For PA, Inferred CNVs included events previously observed including gains on chromosomes 5, 7, 15, and 20 (17, 19). In summary, our scRNA-seq data revealed both neoplastic and non-neoplastic cells with non-neoplastic cells comprised of both myeloid and lymphocytic cell lineages.

### pLGG neoplastic population consists of distinct subpopulations in PA and GG

To assess differences in cellular identity between pLGG subtypes, neoplastic cells were separated from non-neoplastic cells and re-clustered using Harmony. We identified nine PA (cell # = 3531) and seven GG subclusters (cell # = 1519) (Figure 2A-B). The number of cells comprising each neoplastic subcluster are detailed in Supplementary Table1. Following further molecular characterization, we observed a developmental hierarchy among both PA and GG subpopulations. These subpopulations include cancer cells resembling oligodendrocyte progenitor cells (OPC-like), oligodendrocytes (OC-like), astrocytes (AC-like), and specifically for GG, neurons (Neuron-like). Additionally, we identified cells with high levels of MAPK genes specifically in PA. Furthermore, several less-defined subpopulations including cancer cells demonstrating hypoxic condition (in PA) as well as cells with high levels of ribosomal and glycolysis activities (in GG) were identified.

**Figure 2:**
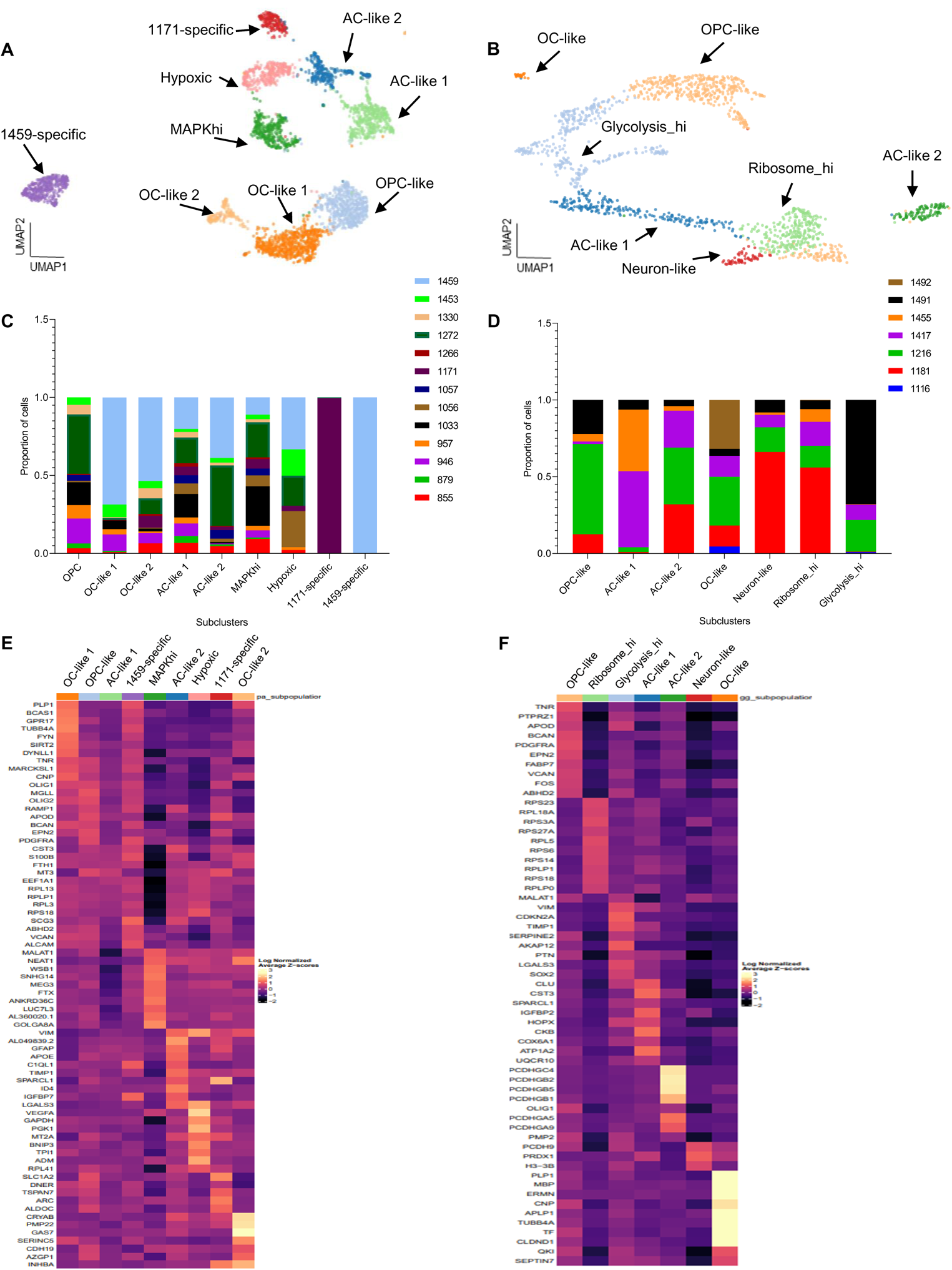
Gene expression profiling of neoplastic populations demonstrate distinct subpopulations in PA and GG. Harmony aligned UMAP plots of **A)** PA cells colored by nine identified subclusters and **B)** GG cells colored by seven identified subclusters. Stacked barcharts show the contribution of each patient sample to various **C)** PA and **D)** GG subclusters. Cell counts for subcluster are normalized to 1. Heatmaps for the top differentially expressed genes in various **E)** PA and **F)** GG subclusters. Each row represents a gene and each column represents a neoplastic cluster. Abbreviations: AC, astrocytes; OC, oligodendrocytes; OPC, oligodendrocyte progenitor cells; MAPKhi, high expression levels of MAPK genes; Glycolysis_hi, high expression levels of genes involved in glycolysis; Ribosome_hi, high expression levels of ribosomal genes.

The most abundant PA subcluster resembled OC (OC-like 1). These cells appear as differentiating oligodendrocytes (Supplementary Figure 3A) differentially expressing genes including *PLP1* as well as *GPR17* and *BCAS1*. Studies have identified *PLP1* and BCAS1 as the most abundant myelin protein and a novel myelin-associated protein, respectively. Interestingly, *BCAS1* has been shown as an OC-like gene signature in a previous study investigating transcriptional landscape of PAs (17, 20). *GPR17* is a suggested regulator of oligodendrocyte development and specification (21). There was a smaller OC-like 2 population, which, although correlated with oligodendrocyte populations within Allen Brain Atlas (Supplementary Figure 4A), appeared to have a unique gene expression profile compared to OC-like 1. CRYAB and PMP22 are among the top differentially expressed genes within OC-like 2 cluster and have shown to be myelin associated proteins (22, 23). Finally in PA we identified a subcluster of cells showing high MAPK activity (Supplementary Figure 5). This subcluster differentially expresses genes enriched in MAPK signaling pathway (MAPKhi) such as WSB1, SNHG14, and MEG3. These long noncoding RNAs have been shown to regulate MAPK pathway (24). The presence of these differentially expressed transcripts in our MAPK subcluster correlate with previous studies by Reitman, Z.J., et al. (17).

An OPC-like population was the largest cell population in GG (Figure 2B) (25). This was one of the shared subclusters between PA and GG. Although in both populations we identified known OPC markers (i.e PDGFR and BCAN), the gene expression patterns differed (Figure 2E-F).

In GG, there was also a small set of OC-like cells resembling a more mature oligodendrocyte population differentially expressing mature OC-related genes such as MBP and ERMN. Astrocyte (AC-like 1-2) subpopulations were identified in both tumor types. Interestingly, all four AC-like populations in PA and GG showed different gene expression patterns (Figure 2E-F). AC-like 1 demonstrated a less-defined gene expression profile relative to other clusters but was correlated with two of the Allen Brain Atlas astrocyte populations (Supplementary Figure 4A); these cells appear to be undergoing a high degree of protein synthesis (Supplementary Figure 3). The smaller AC-like population in PA (AC-like 2), showed differential expression of GFAP and VIM, two known intermediate filaments in astrocytes. AC-like 1 in GG contained several members of COX genes which are involved in mitochondrial respiratory chain complexes metabolism (Supplementary Figure 3). The second less populated astrocytic cluster (AC-like 2) showed differential expression in several genes (PCDHGC4, PCDHGB1, PCDHGB2, PCDHGB5) encoding cell adhesion molecules (Supplementary Figure 3) that regulate astrocyte-neuron interaction. The remaining identified subclusters for each population are detailed in Supplementary Figure 3. Marker genes identifying each PA and GG subcluster are shown in Supplementary Table 2 and 3 respectively.

There were two sample-specific clusters (UPN#1459 and UPN#1171). In GG, there were cell populations that we could not clearly define, one carrying high ribosomal genes (Ribosome_hi) and one with high levels of glycolytic activity (Glycolysis_hi). In summary, we observed several neoplastic subclusters in both PA and GG which revealed a developmental hierarchy demonstrating immature and mature brain cancer cells. Interestingly, even clusters identified having the same origin had different gene expression patterns.

### Characterizing immune populations in PA and GG

To fully characterize immune subclusters, we separated the non-neoplastic subclusters into myeloid and lymphoid lineages. Unsupervised clustering of each using Harmony revealed 10 myeloid lineage and eight lymphoid lineage subclusters (Figure 3A-B). The number of cells comprising each immune subpopulation are detailed in Supplementary Table 4. Marker genes identifying each subcluster are shown in Supplementary Table 5 and 6. We identified three microglia populations differentially expressing P2RY12, a known marker of microglia (Microglia Comp+P2RY12), Microglia P2RY12+, and Microglia CCL3+P2RY12. Microglia CCL3+P2RY12 appeared to be functionally more activated expressing high levels of chemokines such as *CCL3* and *CCL4*. Additionally, we identified a highly chemotactic myeloid subpopulation (Myeloid-Chemokine), which also expressed chemokine molecules *CCL3* and *CCL4*. Among smaller clusters, we identified a myeloid-neuron mix population (Myeloid-Neuron) differentially expressing *MAP1B*, a neurogenesis-related gene, a dendritic cell population (DC-like myeloid), as well as a small neutrophil and M2 macrophage population differentially expressing *FCER1A* and *MRC1*, respectively. We also detected a very small subset of cells differentially expressing genes associated with hypoxia (*MIF* and *LDHA*). All samples contributed to all myeloid subclusters to various degrees (Supplementary Figure 6A). We have mainly used non-conventional identities to annotate myeloid subclusters, as myeloid and lymphoid cells shift away from the conventional identities in the presence of tumor microenvironment.

**Figure 3:**
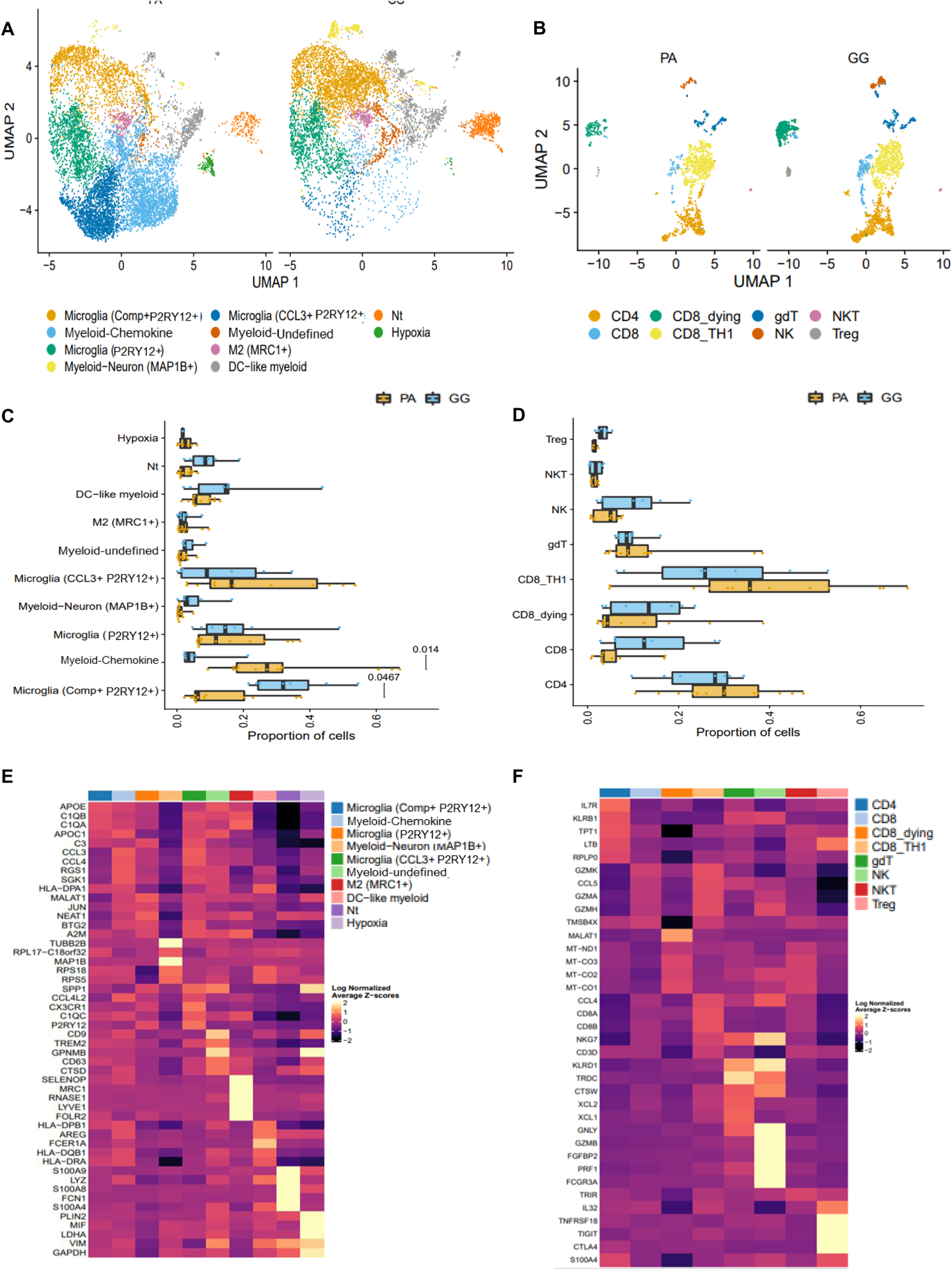
scRNA-seq analyses of immune cells in PA and GG. Harmony aligned UMAP plots of **A)** myeloid and **B)** T cell subclusters. Bargraphs comparing cell proportions in **C)** myeloid and **D)** T cell populations between PA and GG. Heatmaps for the top differentially expressed genes in various **E)** myeloid and **F)** T cell subclusters. Each row represents a gene and each column represents a non-neoplastic cluster. Abbreviations: Nt, neutrophils; DC, dendritic cells; Treg, regulatory T cells; NKT, natural killer T cells; NK, natural killer cells; gdt, gamma delta T cells; PA, pilocytic astrocytoma; GG, ganglioglioma.

We compared the proportion of cells in myeloid subclusters between PA and GG samples (Figure 3C-D). While there appeared to be differences in cell proportions among all myeloid subclusters, we noticed a significant difference in cell proportion particularly in two subclusters. Cells comprising the Myeloid-Chemokine cluster were significantly more prevalent in PA compared to GG (p=0.014). In comparison, Microglia (Comp+ P2RY12+) had a significantly higher prevalence in GG (p=0.0467).

Unlike in the myeloid subclusters there was no significant difference among T cell subclusters between PA and GG (Figure 3D). All patient samples contributed to the different T subclusters (Supplementary Figure 6B). Harmony clustering identified several T cell subclusters (Figure 3B) as well as smaller subsets of NK and NKT cells. The top differentially expressed genes in each non-neoplastic subcluster is shown (Figure 3 E-F).

Our data revealed several myeloid and T cell subclusters that contributed to PA and GG tumor microenvironments. Interestingly, a significant difference was seen between the prevalence of major myeloid subclusters between PA and GG. To gain a better understanding of this underlying difference, we decided to study the immune signals via which these cells communicate by analyzing their secreted cytokines.

### KIAA-1549:BRAF fusion tumors appear to have higher levels of immune mobilizing cytokines compared to BRAF V600E tumors

scRNA sequencing analysis of our immune populations led us to hypothesize that there would be differences in the cytokine signaling from myeloid cell populations between PAs and GGs and across mutational profiles (*KIAA-1549:BRAF* fusion, *BRAF* V600E, *FGFR3*). To investigate this, we performed a broad cytokine analysis utilizing two platforms to gain a better understanding of the function that each myeloid cell subpopulation has within pLGG. We performed bulk cytokine analysis on media collected from tumor and immune cells following a short culture and compared the secretion profile between *BRAF* V600E and *KIAA-1549:BRAF* fusion samples. Though not significant, there was a trend towards higher concentrations of cytokines secreted by KIAA-1549:BRAF fusion samples for both tumor and immune cells of (Figure 4A-B, Supplementary Figure 7).

**Figure 4:**
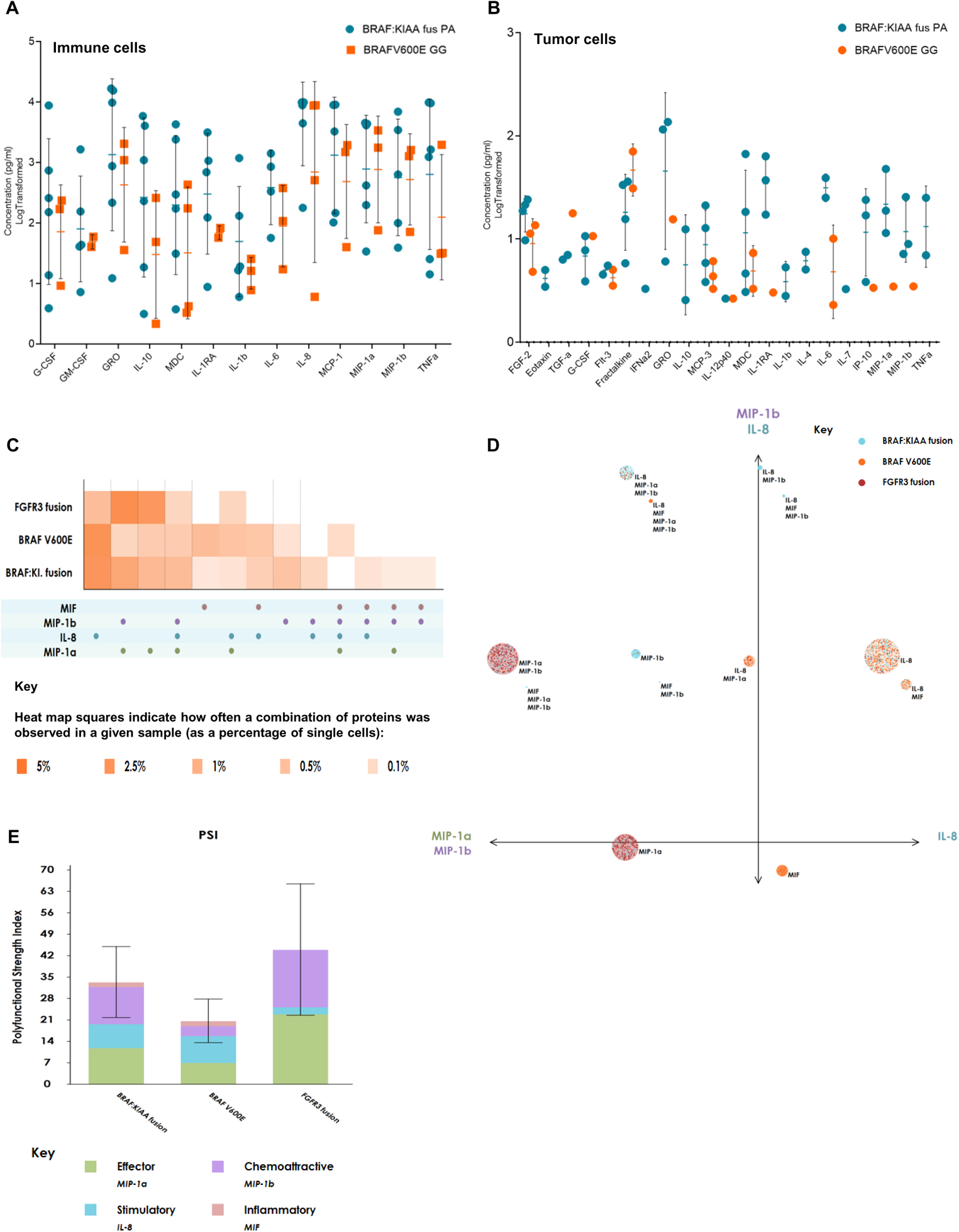
Bulk and single cell cytokine/chemokine analyses in PA and GG. Dot plots of detected cytokines/chemokines from bulk **A)** myeloid (cytokines/chemokines with concentrations greater than 1000pg/ml) and **B)** tumor (all cytokines/chemokines) cells conditioned media in PA and GG. **C)** Single cell polyfunctional heatmap displaying functional cytokines/chemokines of interest secreted across myeloid cells from samples PA and GG samples with different mutational status. The heatmap is color-coded from light to dark, based on the frequency of the polyfunctional subsets. The rows of squares correspond to the three sample groups and the columns correspond to polyfunctional group of cytokines that was expressed in at least one of the sample groups. **D)** Polyfunctional Activity Topography Principle Component Analysis (PAT PCA) of particular functional groups across samples with different mutational status. Each color coded dot represented a single cell from one of our three sample groups. The size of the circles corresponds to the frequency of an individual polyfunctional group (large groups = large circles, small groups = small circles), and the cytokines labeled next to it represent the cytokines present in the functional group The principle components are labeled according to their correlation with specific cytokines. PC1 and PC2 are a combination of dominant cytokines that drive the polyfunctionality of myeloid cells. **E)** Polyfunctional Strength Index (PSI) computed for myeloid cells at the single cell level across three sample groups with different mutational status.

To get a better understanding of the cytokine signaling in PA and GG myeloid cells, we performed single cell cytokine analyses. To distinguish distinct polyfunctional myeloid subsets and the heterogeneity within our patient samples, we utilized a polyfunctional heatmap (Figure 4C). This visualization shows major functional subsets secreted across different mutational groups (*KIAA-1549:BRAF* fusion, *BRAF* V600E, and *FGFR3* fusion samples). As seen in Figure 4C, there is heterogeneity in polyfunctional groups across all sample groups. Overall, *KIAA-1549:BRAF* fusion samples appear to have the most polyfunctional groups. It is interesting to note that there is a higher percentage of single cells in the *KIAA-1549:BRAF* fusion group compared to the *BRAF V600E* group that express a combination of chemokines, MIP-1α (CCL3) and MIP-1β (CCL4) as well as a combination of MIP-1α, MIP-1β, and IL-8 (CXCL8). A higher percentage of single cells in the *BRAF V600E* group express the pro-tumor cytokine MIF, but this expression pattern changes when MIF is expressed along with the above-mentioned chemokines.

To visualize distinct polyfunctional myeloid subsets and their complex landscape, we performed a modified PCA analysis called Polyfunctional Activation Topology Principle Component Analysis (PAT PCA). In the resulting scatter plot (Figure 4D), each functional group discussed in the heatmap above is represented by a circle in the PAT PCA graph. A number of functional groups are identified, each secreting a particular combination of cytokines. Consistent with what we identified in the polyfunctional heatmap, the PAT PCA graph also shows higher levels of MIP-1α and MIP-1β dominated polyfunctional groups secreted by BRAF and FGFR3 fusion samples compared to *BRAF V600E* single cells. Our data show that there are distinct populations of cells that are secreting one or more cytokines/chemokines across mutational subgroups. BRAF fusion cells appear to be in a more immunologically active state. In summary, these data highlight a functional difference between *KIAA-1549: BRAF* fusion and *BRAF V600E* immune cells.

Finally, to quantify the collective impact of polyfunctional cells (simultaneously secreting multiple cytokines at high intensity), we used a poly functional strength index (PSI). PSI is defined as the percentage of polyfunctional cells within a particular sample multiplied by the average signal intensity of the cytokines that are secreted from those cells. The PSI revealed that chemokines including MIP-1α and MIP-1β have a higher impact in BRAF fusion tumors compared to *BRAF V600E* tumors (Figure 4E). Collectively, these data suggest a complex and distinct immune phenotype between *KIAA1549-BRAF* fusion and *BRAF V600E* suggesting the involvement of potentially different immune pathways in these tumor subtypes.

### Spatial transcriptomics (ST) reveal differential expression of chemokines in PA and GG

Both scRNA-seq and single cell cytokine assays, revealed differences in immune cell infiltration and function between PA and GG. However, single cell techniques lack spatial orientation of cells within the TME. Therefore, we performed ST analysis in both PA and GG using the Visium platform (10X genomics). This analysis was performed on five PA (UPN#946, UPN#1056, UPN#1272, UPN#1330, UPN#1453) and three GG (UPN#1181, UPN#1491, UPN#1492) snap frozen surgical sections. All PA and GG harbored BRAF fusion or BRAF V600E alterations respectively. ST data was processed and filtered resulting in (total number of spots =14,990 (10,477 spots per PA and 4,513 per GG samples)) across all samples. Sample spots were clustered with batch correction using harmony resulting in five PA and three GG clusters across the eight samples (Figure 5A-B).

**Figure 5:**
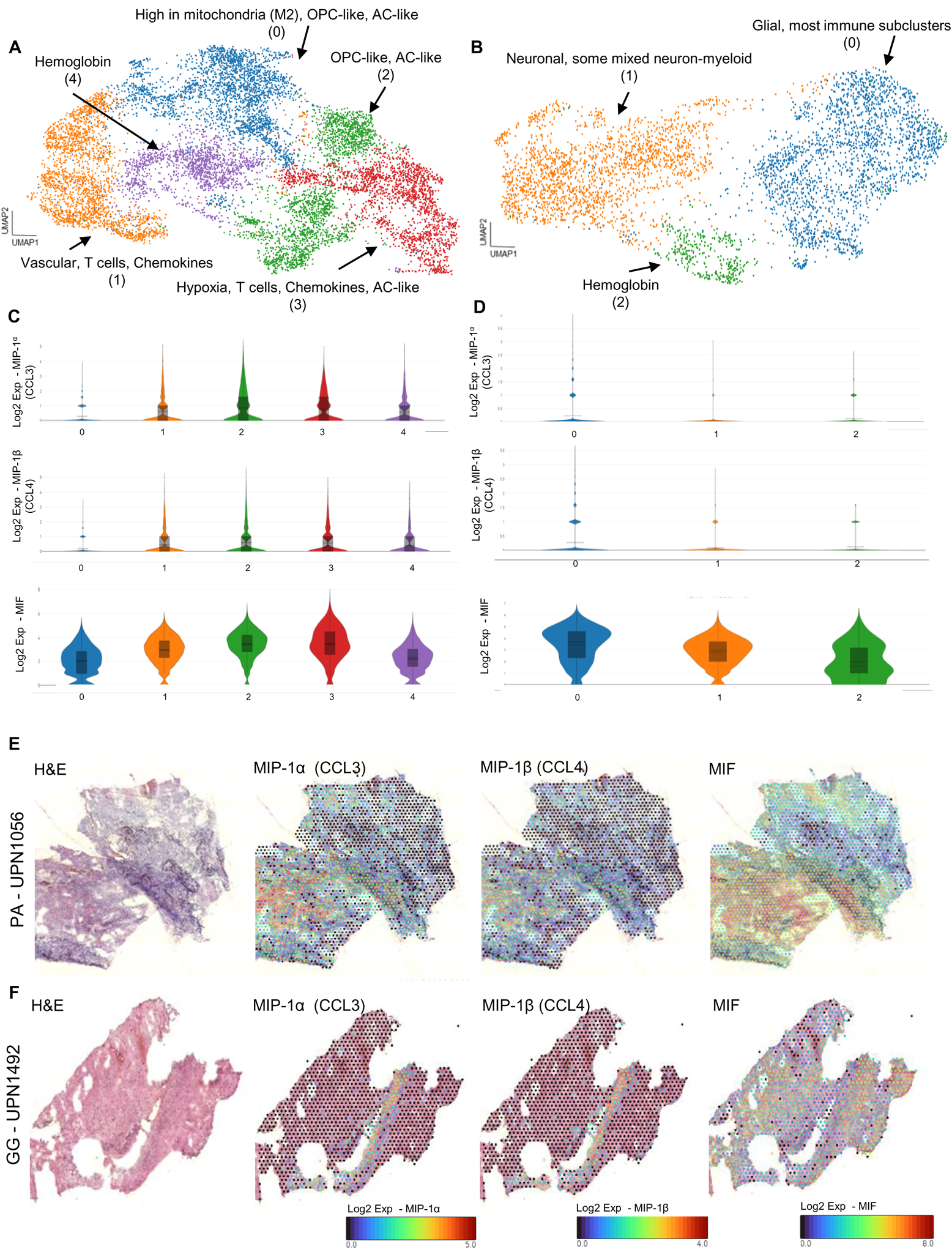
cytokines/chemokines have a differential distribution within TME in PA and GG. UMAP clustering of spatial transcriptomic data colored by different subclusters in **A)** PA and **B)** GG. Each subcluster comprises of different neoplastic and non-neoplastic cell types. Violin plots showing the expression levels of MIP-1α, MIP-1β, and MIF in **C)** PA and **D)** GG. Representative images showing spatial expression levels of MIP-1α, MIP-1β, and MIF in a **E)** PA and **F)** GG sample.

Transcriptomic profiles from each spot cluster were examined using clustree analysis of ST cluster similarity and ontological analysis of ST marker genes. Different clustering resolutions were utilized for identifying cell types present in various regions of clusters. ST showed distinct regions within PA and GG tumors consisting of both immune and tumor cell populations (Figure 5A-B). Marker genes identifying cell types within various regions of PA and GG are shown in Supplementary Table 7 and 8 respectively. As predicted by our cytokine data, higher expression of genes associated with identified cytokines (MIB-1α, MIP-1β, and MIF) was observed in PA compared to GG samples (Figure 5C-F, Supplementary Figure 8). These data support our cytokine analysis, indicating an importance and difference in cytokine gene expression between pLGG groups.

## Discussion

pLGG are a heterogeneous set of tumors with varying clinical behavior and varying responses to therapy. To help gain a deeper understanding of the heterogeneity of the tumor microenvironment and the potential role immune cells may play in pLGG, we used a three pronged approach with (1) scRNA-seq to biologically characterize two of the most common pLGG subtypes PA and GG (2) bulk and single cell cytokine analysis for a more functional understanding of these tumors, and (3) ST analyses as a validation method for findings from our scRNA-seq and cytokine analyses. We believe such a multi-pronged approach comparing different tumor subtypes will be needed to fully understand how to develop successful immunotherapy for pLGG.

Previous data evaluating small populations of PA and GG assessed the tumor types independently (17, 18). Our data allowed us to compare and contrast tumor subpopulations among our PA and GG tumor samples. Our results demonstrate a developmental hierarchy with immature progenitor cells and mature oligodendrocytes and astrocytes. When compared with the previous PA study, our MAPKhi population showed the strongest correlation with their MAPKhi population. There was no substantial overlap between our OC-like and AC-like populations and theirs. This could be due to different methods of scRNA seq analysis resulting in differential capture of markers and limited sample size. Analysis of GG tumors using single nuclei RNA seq provided a comprehensive review of the transcriptomic landscape of these samples and a potential prognostic signature for this tumor type (18). Their data would support our findings that GG may be less immunogenic and therefore less responsive to immune therapies.

Our scRNA-seq analyses revealed several immune populations. The most notable observation was identification of a large myeloid subpopulation in our overall pLGG dataset, comprising over 50% of cells. We identified several myeloid subclusters and observed a significant difference in two distinct subclusters (chemokine-myeloid and microglia (comp+ P2RY12+)) between PA and GG. There were a significantly higher number of cells comprising chemokine-myeloid subcluster in PA compared to GG and a larger microglia (comp+ P2RY12+) subcluster in GG. The chemokine-myeloid subcluster demonstrated an increased expression of genes responsible for the immune mobilizing chemokines MIP-1α and MIP-1β. Since chemokines are originally known to recruit other immune cells to the tissue site during both homeostasis and in response to infection and inflammation, this observation may suggest a higher level of immune activity in PA compared to GG.

Our scRNA seq data suggested cytokine signaling may be important to the overall immune function of these tumors. To evaluate the potential cytokine differences and examine the level of immune activity in our PA and GG samples, we performed bulk and single cell cytokine analyses. We observed specific chemokines such as MIP-1α and MIP-1β to be secreted in higher concentrations in fusion samples compared to BRAF V600E samples. This is in agreement with our scRNA-seq analysis data. It has been shown that MIP-1α plays a critical role in recruiting distinct immune phenotypes to intratumoral sites and it induces antigen specific T cell responses (26). Our work has implications for vaccine trials aiming to utilize the power of the immune system against pLGG (27). The functional cytokine differences identified in our data suggest immunotherapy approaches towards tumors with *KIAA1548-BRAF* fusion and *BRAF V600E* mutations may need to be customized for each group to capitalize and compensate for innate cytokine differences between them. Future studies will investigate the cytokine profiles from T cell populations isolated from our patient samples to better understand the functional classes of T lymphocytes, including exhausted T cells, which may help us assess the downstream impact of elevated chemokines and cytokines in PA tumors. To see if we can observe the differential expression of chemokines and cytokines within the tumor cellular architecture in intact tissues, we utilized spatial transcriptomics. Our data showed higher expression of chemokines of interest in the spatial environment of PA tumors compared to GG tumors. This was consistent with our scRNA seq and the cytokine data.

As with all studies, our analysis carries some limitations. A larger sample size (especially for GG subtype) resulting in higher number of cells could provide a clearer picture of the cellular identity of our immune and tumor subclusters. Additionally, future studies of lineage analyses will be needed for a deeper understanding of the identified subclusters. Additional future studies will also focus on how not only genetic drives of tumors, but also how location of tumors may influence these results. Finally, future additional pathway and functional analysis will be performed to better understand the downstream impact of elevated chemokines and cytokines in our PA fusion samples and to gain a broader understanding of the biology behind and affected by these processes.

Despite some of these limitations, we believe our study has expanded the critically important genomic and functional understanding needed for the role immune cells may play in the TME in the context of pLGG. Data has revealed that pLGG contains a vast immune population, but there are differences between tumor types and genetic changes that may influence their response to immunotherapy. Future proteomic studies are ongoing to complement our transcriptomic analyses. This comprehensive approach helps further our understanding of the potential impact of molecular differences on tumor biology and capitalize on the potential of immunotherapies for pLGG patients.

Further investigation of the immune cell infiltration and tumor-immune interactions is warranted.

## Material and Methods

### Sex as a biological variable

Our study examined male and female patient samples and similar findings are reported for both sexes.

### Patient sample acquisition and preparation

Human pLGG samples (n=23) were collected from surgeries at Children’s Hospital Colorado with IRB approval (COM-IRB 95-500) (Table 1). For single cell approaches, tissue was collected in serum-free media, brought to the laboratory for mechanical dissociation, and viably frozen in single cell suspension as previously described (28). Samples utilized for spatial transcriptomics and cytokine analyses were snap frozen at surgery.

### ScRNA-seq analysis

scRNA-seq was performed on 23 pLGG patient samples. Prior to sequencing, samples were thawed in batches and flow sorted (Astrios EQ) to obtain viable single cells based on <u>propidium iodide</u> (PI) exclusion. With the study goal of performing scRNA-seq on 2,000 cells per sample, we utilized a Chromium Controller in combination with Chromium Single Cell V3 Chemistry Library Kits, Gel Bead & Multiplex Kit and Chip Kit (10X Genomics). Single cells were isolated into microfluidic droplets containing oligonucleotide-covered gel beads that capture and barcode the transcripts. Transcripts were converted to cDNA, barcoded and sequenced on Illumina NovaSeq 6000 sequencer to obtain approximately 50 thousand reads per cell.

### scRNA-seq data analysis

Raw sequencing reads were processed into gene-expression matrices using Alevin (salmon version 1.3.0) with the hg38 human reference genome. The resulting count matrices were further filtered in Seurat (v.4.0.3-4.1.1) (https://satijalab.org/seurat/) to remove cell barcodes with less than 200 genes, more than 20% of UMIs derived from <u>mitochondrial genes</u>, or less than 500 UMIs and more than 50,000 UMIs. UMI counts were log-normalized. PCA was performed on scaled normalized counts using highly variable genes (2,000). Harmony (v0.1.0) was applied to integrate samples (theta = 1). 25 harmony components were used for generating a UMAP projection and a shared nearest neighbor graph, which was clustered using graph based clustering implemented in Seurat. Doublet cells were identified using scDblFinder, and cells with doublet scores greater than 5 were excluded (v. 1.6.0). Cells were classified into broad cell types (T/NK, B, Myeloid, Neutrophil, Glioma, and Proliferating) by comparing clusters to a single cell reference dataset using clustifyr (v1.4.0). The custom reference was constructed using single cell RNA-seq from other pediatric brain tumors and normal human cell datasets (20, 28–32). Tumor cells from individual subgroups, and immune cell types (T and Myeloid) were reprocessed separately for further analysis. Differential expression and marker gene identification was performed using findMarkers() from the scran package (v. 1.20.1), with the block argument set to each sample to control for differences in sample compositions in each cluster. The presence of copy number variants (CNVs) was identified using inferCNV.

### Cytokine*/*Chemokine analysis

Patient samples were thawed and incubated at 37°C for 24 hours. Using CD45, myeloid and tumor cells were separated and incubated at 37°C for an additional 24 hours. LPS stimulation was performed on the myeloid cells. The condition media was then collected and stored in −80 freezer for cytokine and chemokine analysis using the Milliplex platform (n=11) and the immune cells were used for single-cell cytokine profiling using the single-cell secretome platform (n=12) on the IsoSpark system (IsoPlexis, Branford, CT).

### Multiplex cytokine bead array assay

Cytokine/chemokine concentrations from bulk conditioned media were measured using Milliplex, following manufacturer’s instructions as previously described (33).

### Single-cell multiplex cytokine and chemokine profiling

Following LPS stimulation, immune cells were collected and prepared for single-cell cytokine capture on Innate Immune IsoCode chips per manufacturer recommendation (IsoPlexis, New Heaven, CN). Data was analyzed using IsoSpark software.

### Spatial transcriptomic analysis

Frozen samples (n=8) were OCT embedded and sectioned at 10μm on a Cryostar NX70 cryostat (Thermo Fisher Scientific). Capture sections were fixed with methanol, stained with H&E, and imaged on an Evos M7000 (ThermoFisher) with brightfield settings. Following image capture, capture sections were permeabilized and processed to generate RNA libraries following 10x Visium protocol. Libraries were sequenced to a depth of 70,000 read pairs per spot calculated from the image, on a Novaseq6000 (Illumina) sequencer.

### Spatial transcriptomics data analysis

Sequencing data were processed with Space Ranger (10x genomics, v1.2.1), followed by further analysis in R using the Seurat tool suite. Spots were filtered to ensure the number of genes detected between 50 and 15000, and less than 50% of UMIs mapped to mitochondrial genes. After initial SCTransform normalization on each sample and principal component analysis on merged data, sample integration was performed with Harmony (v0.1.0) using 30 principal components. UMAP dimension reduction and shared nearest neighbor clustering were carried out on 30 principal components, and clustering results at different resolution settings were explored through Clustree (v0.4.3) visualizations.

### Deconvolution

Bulk tumor tissue analyzed via Affymetrix array (n=36) was imputed into CYBERSORTx to further explore the single-cell sequencing findings. This dataset contained 7 GG, 22 PA and 6 LGG. A CYBERSORTx signature file was generated based on the pLGG scRNA seq data (<u>GSE232316</u>). P-values for the bulk gene expression datasets and the single-cell derived signature file had a median of 0 (Range 0-0.4). CYBERSORTx analyses were performed in R on absolute mode with 100 permutations using normalized, but not log converted, counts (Mas5.0 normalization for expression array, TPM normalization for RNA-seq).

### Statistical analysis

All statistical analyses were performed using R and Prism (GraphPad). For all tests, statistical significance was defined as P-value < 0.05.

### Study approval

Human samples were collected from surgeries at Children’s Hospital Colorado with IRB approval (COM-IRB 95-500) (Table 1). Written informed consent was received prior to participation.

### Data and code availability

ScRNA-seq and ST data have been deposited in the National Center for Biotechnology Information Gene Expression Omnibus (GEO) database and are publicly accessible through GEO SuperSeries accession number <u>GSE232316</u> (https://www.ncbi.nlm.nih.gov/geo/query/acc.cgi?acc=GSE232316). Analysis code will be provided in a github repository upon publication.

## Authorship

SZ and JML designed the project; KR and RF performed bioinformatics analyses; SZ wrote the initial manuscript; SZ, AD, and AG prepared samples for analyses; AD and AG assisted with design and interpretation of spatial transcriptomic and cytokine analyses respectively; TH and MR provided samples; KR, RF, SZ, AG, AD prepared and reviewed figures; JML, MC, JD, JR, MG, EB, AG, AD, RV, NW, NF, KR participated in data and manuscript review and edit.

## Acknowledgements

The Anschutz Medical Campus Genomics Core performed single cell RNA sequencing and spatial transcriptomic analysis.

