## Supplementary Figures for "Multi-pronged analysis of pediatric low-grade glioma reveals a unique tumor microenvironment associated with BRAF alterations"

### Supplementary Figure 1

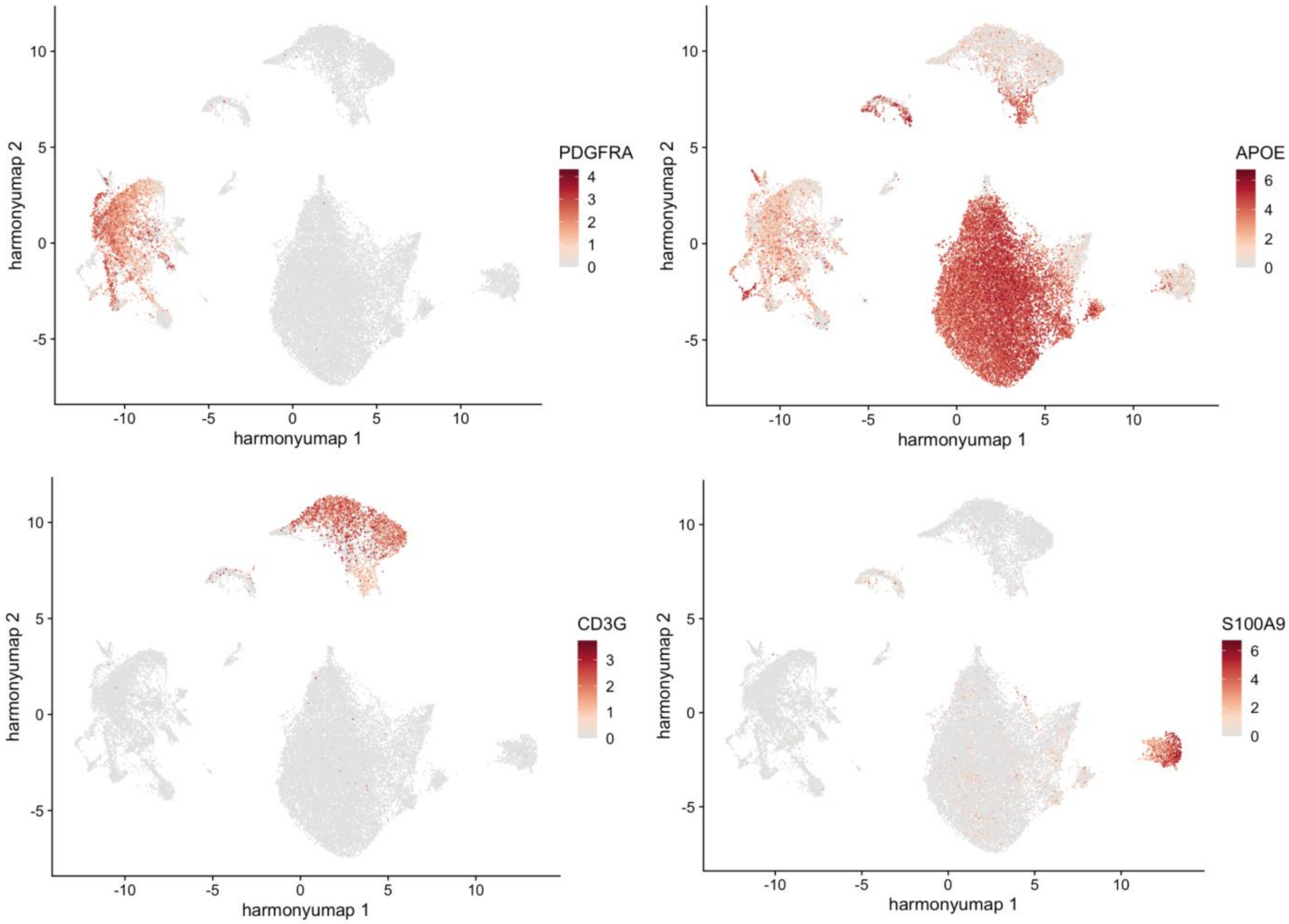


**Supplementary Figure 1.** Feature plots for a neoplastic marker (PDGFRA) and non-neoplastic markers (APOE for myeloid, CD3G for T cell, and S100A9 for Nt). Expression is represented by color gradient with high expression represented by brown and low expression represented by grey.
Abbreviations: Nt, neutrophils.

### Supplementary Figure 2

CIBERSORTx: Using scRNA-seq data to Deconvolve cell subsets from bulk RNAseq data

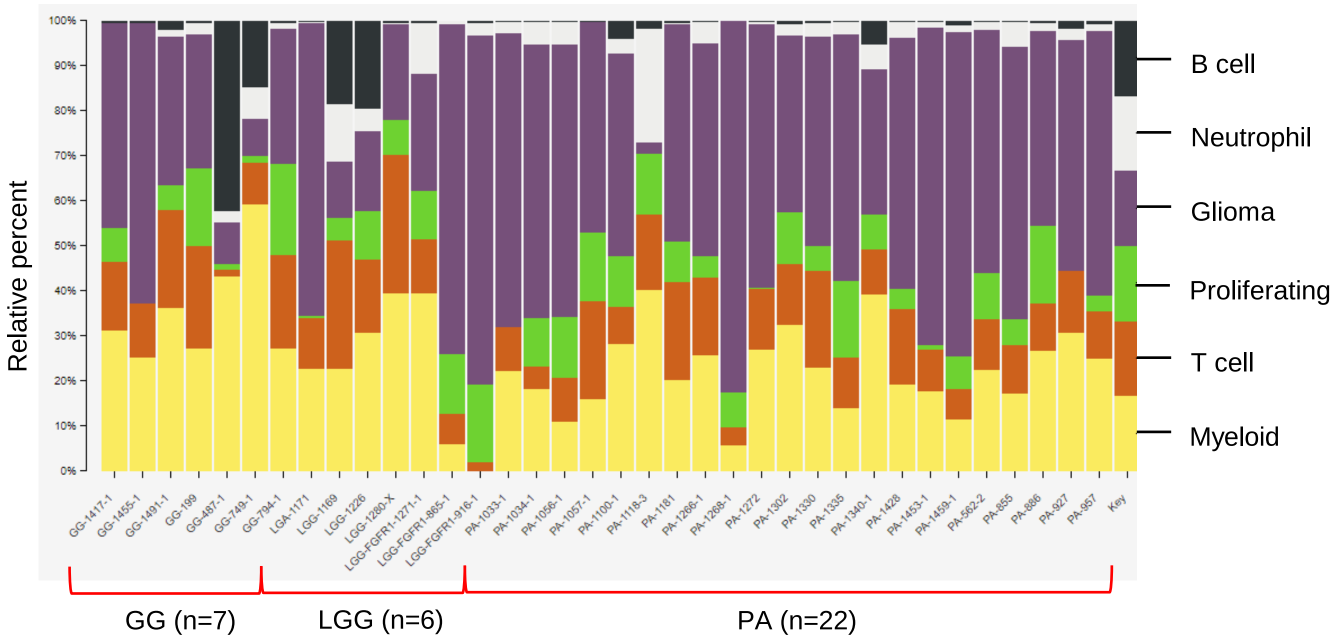


**Supplementary Figure 2.** Deconvolution of cell subsets from pLGG bulk RNAseq data.

### Supplementary Figure 3

**A**

**OC-like 1**

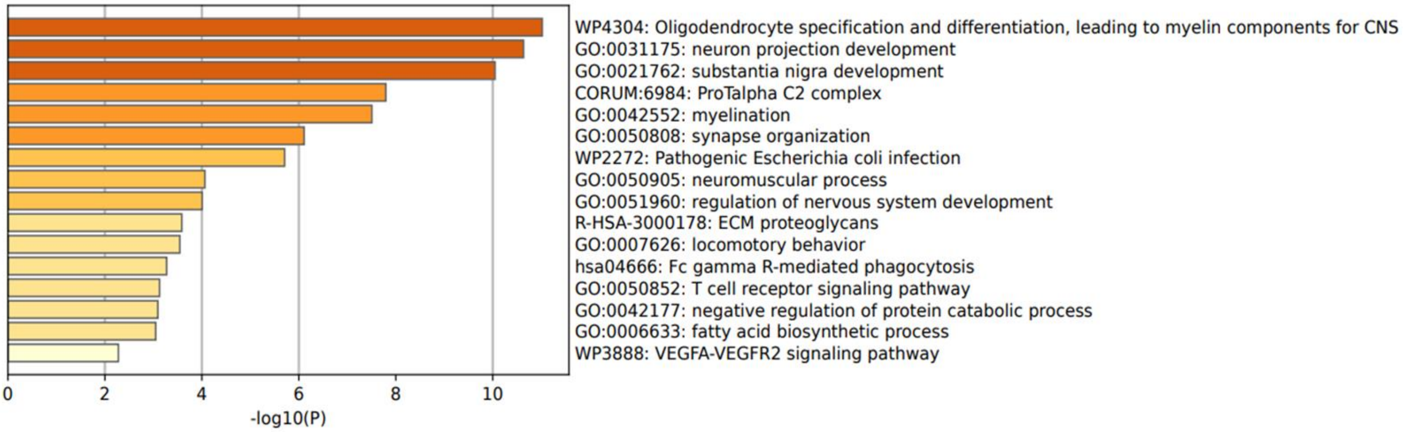


**OPC-like**

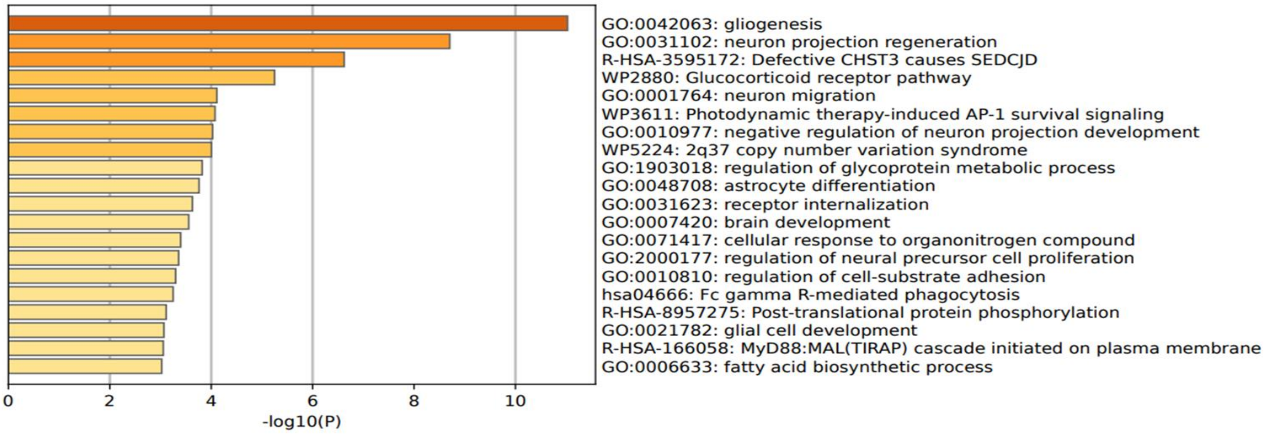


**AC-like 1**

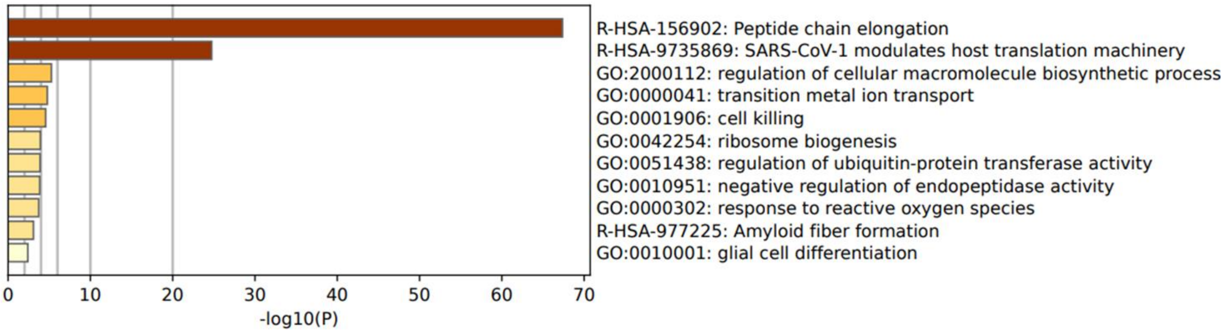


**MAPKhi**

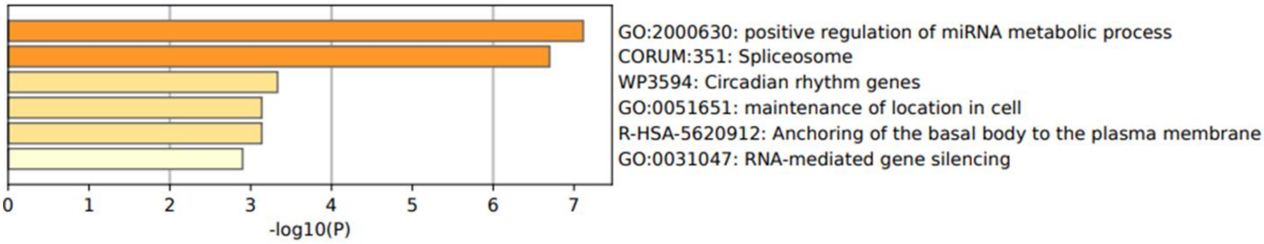


*Note: enrichment results obtained by utilizing all 59 genes

#### Supplementary Figure 3 - continued

**AC-like 2**

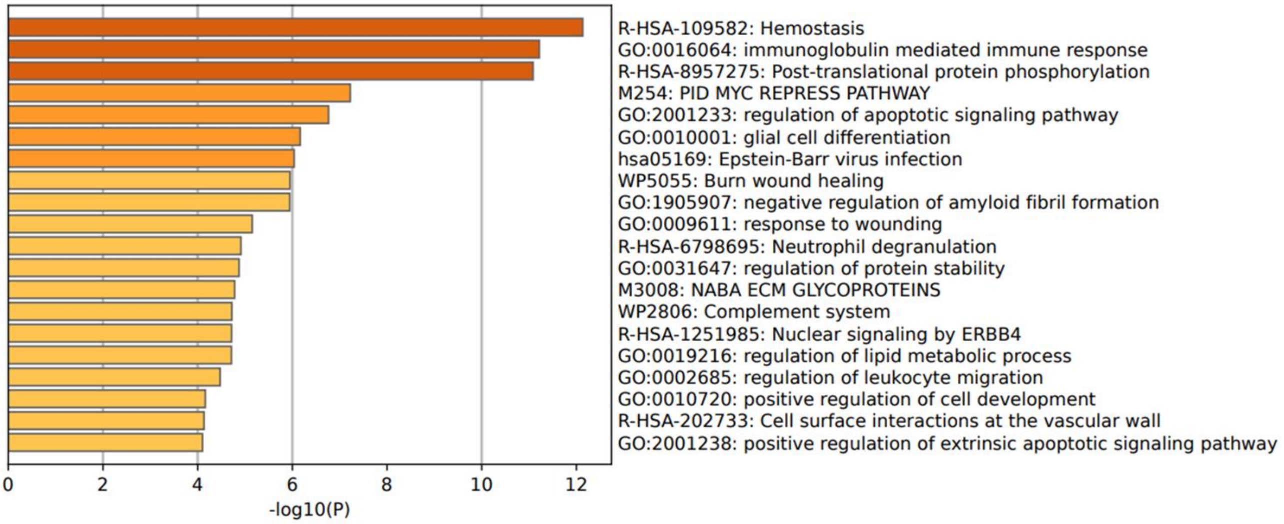


**Hypoxic**

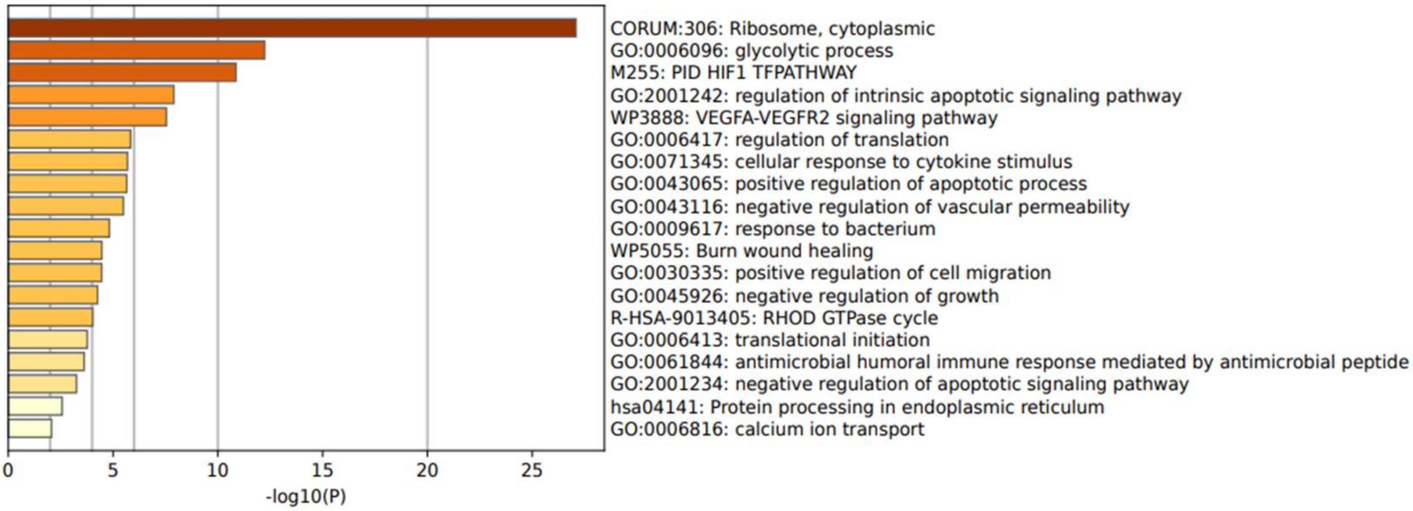


**OC-like 2**

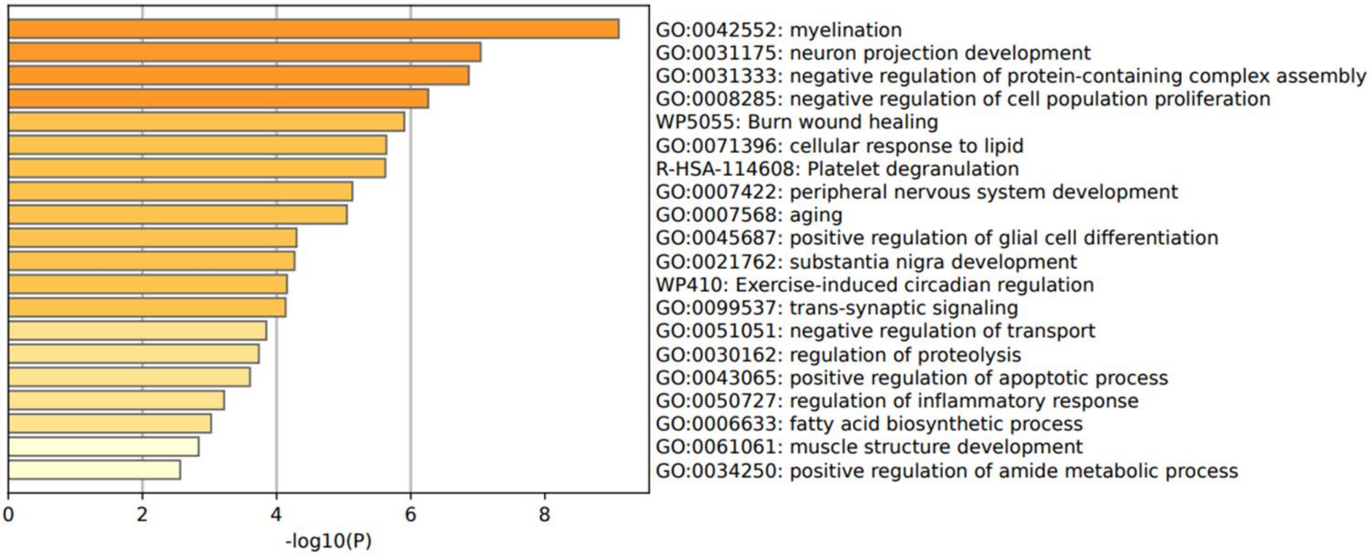

#### Supplementary Figure 3- continued

**B**

**OPC-like**

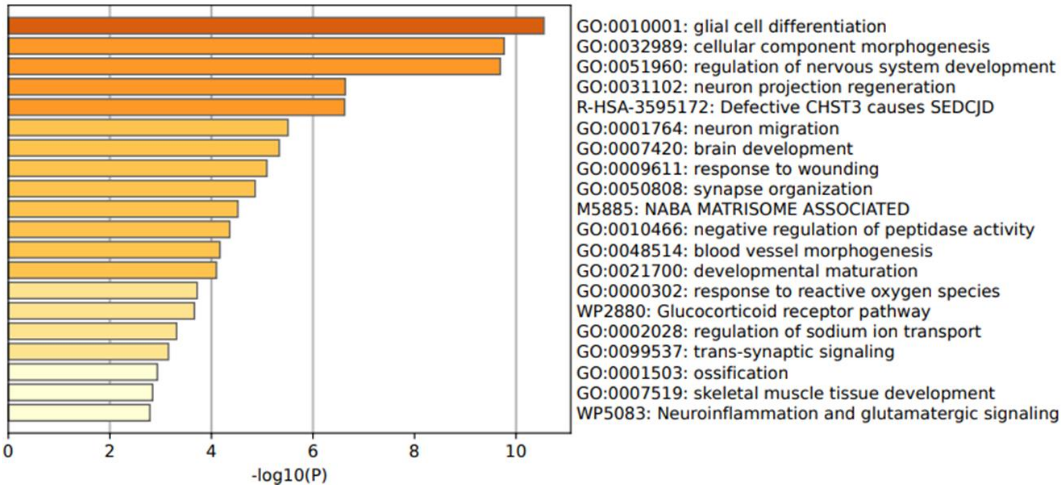


**Glycolysis_hi**

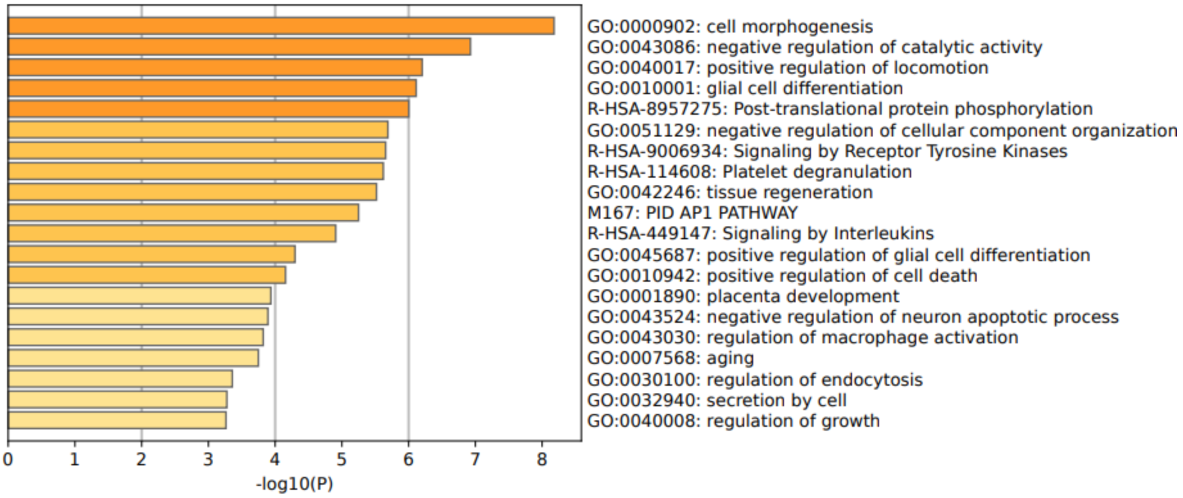


**Ribosome_hi**

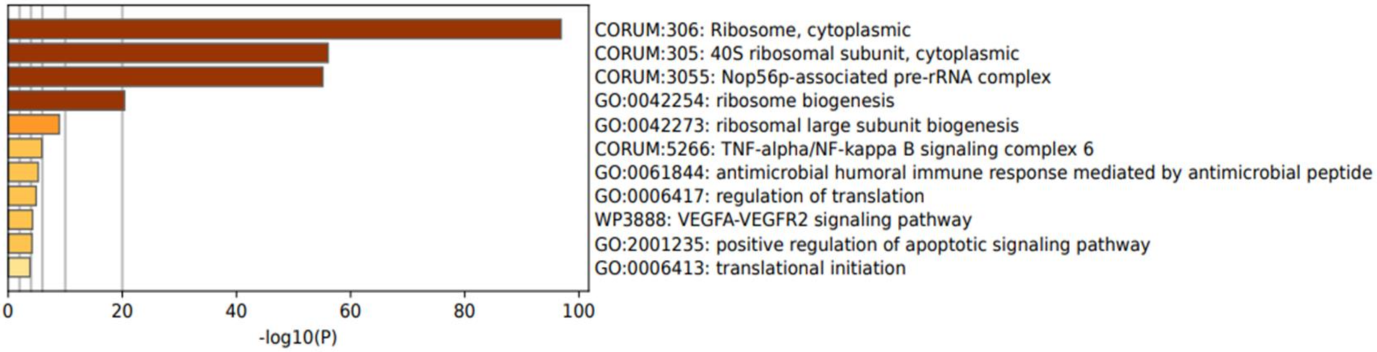


*Note: enrichment results obtained by utilizing all 48 genes

#### Supplementary Figure 3- continued

##### AC-like 1

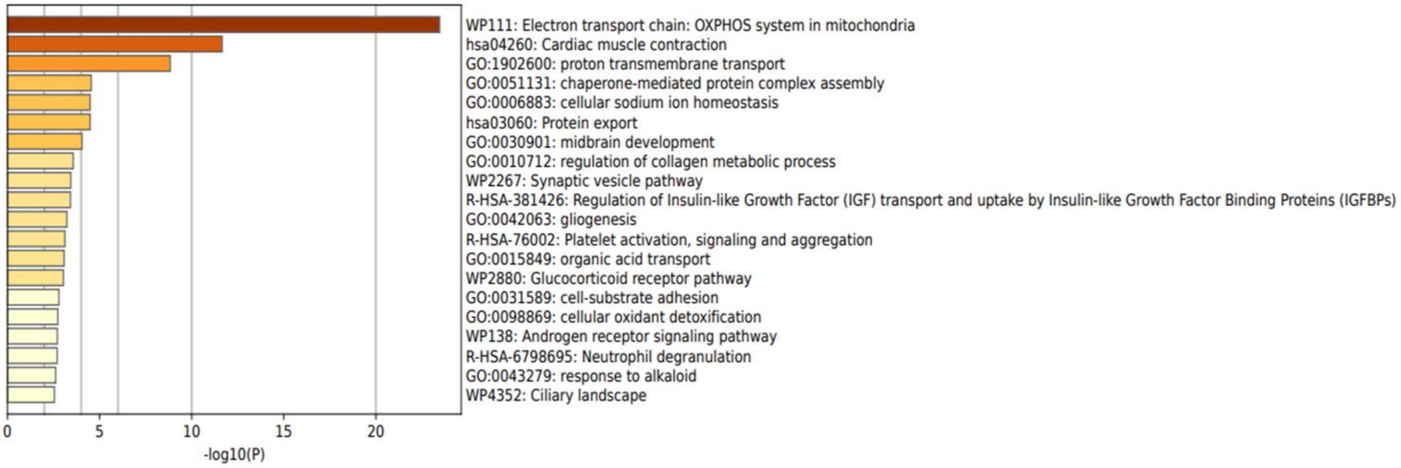


##### AC-like 2

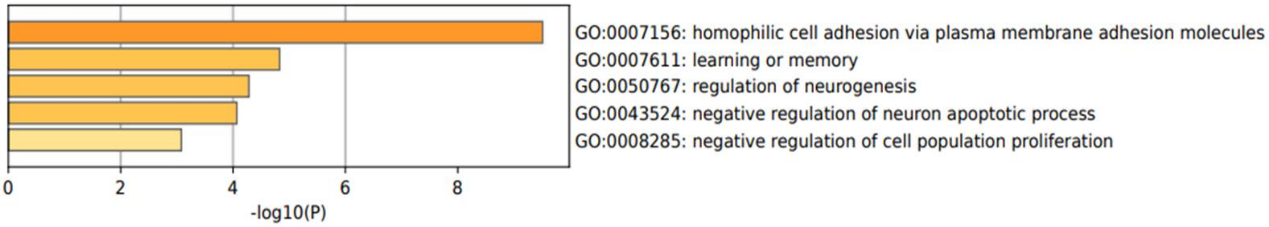


##### Neuron-like

*Note: Only 4 genes were detected following gene expression analysis. Metascape provided no output.

##### OC-like

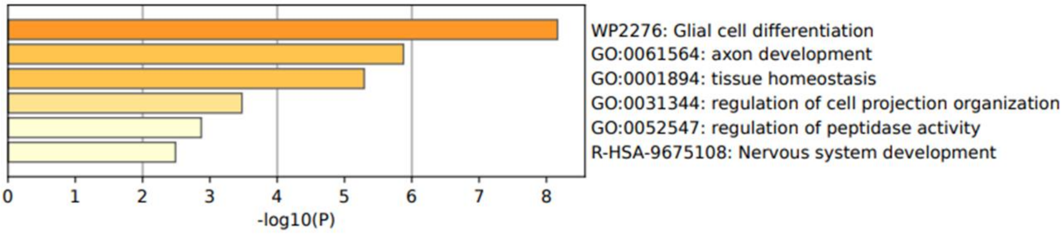


**Supplementary Figure 3.** The cosine similarity between the gene expression values in the single cell data and the average expression data in the data from the Allen Brain Atlas (aba) was computed by using the top 1000 variable genes. Metascape bar graphs using the top 50 differentially expressed genes in each subcluster for viewing top non-redundant enrichment clusters in **A)** PA, **B** GG, one per cluster, using a discrete color scale to represent statistical significance.

### Supplementary Figure 4

**A**

| pa_cell_type | OPC_aba | OC_aba 1 | OC_aba 2 | AC_aba 1 | AC_aba 2 | AC_aba 3 | Exc_aba 1 | Exc_aba 2 | Exc_aba 3 | Exc_aba 4 | Exc_aba 5 | Exc_aba 6 | Inh_aba |
|---|---|---|---|---|---|---|---|---|---|---|---|---|---|
| OPC-like | 0.367 | 0.274 | 0.248 | 0.404 | 0.371 | 0.314 | 0.374 | 0.344 | 0.349 | 0.334 | 0.335 | 0.336 | 0.346 |
| OC-like 1 | 0.392 | 0.421 | 0.394 | 0.335 | 0.312 | 0.272 | 0.411 | 0.366 | 0.390 | 0.356 | 0.367 | 0.371 | 0.373 |
| OC-like 2 | 0.300 | 0.457 | 0.420 | 0.403 | 0.349 | 0.299 | 0.369 | 0.346 | 0.359 | 0.331 | 0.343 | 0.343 | 0.337 |
| AC-like 1 | 0.358 | 0.329 | 0.297 | 0.438 | 0.410 | 0.344 | 0.401 | 0.372 | 0.380 | 0.361 | 0.360 | 0.360 | 0.371 |
| AC-like 2 | 0.255 | 0.257 | 0.228 | 0.455 | 0.415 | 0.345 | 0.348 | 0.332 | 0.332 | 0.321 | 0.320 | 0.318 | 0.326 |
| MAPKhi | 0.365 | 0.301 | 0.269 | 0.424 | 0.379 | 0.350 | 0.358 | 0.334 | 0.343 | 0.321 | 0.364 | 0.360 | 0.326 |
| Hypoxic | 0.216 | 0.244 | 0.210 | 0.368 | 0.333 | 0.284 | 0.378 | 0.363 | 0.349 | 0.353 | 0.345 | 0.342 | 0.346 |
| 1171-specific | 0.362 | 0.242 | 0.209 | 0.444 | 0.418 | 0.359 | 0.397 | 0.378 | 0.375 | 0.368 | 0.367 | 0.366 | 0.362 |
| 1459-specific | 0.336 | 0.344 | 0.316 | 0.347 | 0.328 | 0.275 | 0.365 | 0.337 | 0.347 | 0.330 | 0.331 | 0.330 | 0.342 |

**B**

| gg_cell_type | OPC_aba | OC_aba 1 | OC_aba 2 | AC_aba 1 | AC_aba 2 | AC_aba 3 | Exc_aba 1 | Exc_aba 2 | Exc_aba 3 | Exc_aba 4 | Inh_aba 1 | Inh_aba 2 | Inh_aba 3 | Inh_aba 4 |
|---|---|---|---|---|---|---|---|---|---|---|---|---|---|---|
| OPC-like | 0.421 | 0.265 | 0.247 | 0.369 | 0.405 | 0.320 | 0.348 | 0.354 | 0.341 | 0.349 | 0.320 | 0.336 | 0.350 | 0.324 |
| OC-like | 0.253 | 0.725 | 0.716 | 0.287 | 0.338 | 0.264 | 0.297 | 0.315 | 0.313 | 0.298 | 0.291 | 0.281 | 0.298 | 0.299 |
| Ribosome_hi | 0.235 | 0.228 | 0.214 | 0.381 | 0.412 | 0.344 | 0.341 | 0.348 | 0.325 | 0.330 | 0.323 | 0.343 | 0.342 | 0.316 |
| AC-like 1 | 0.213 | 0.197 | 0.177 | 0.519 | 0.518 | 0.493 | 0.329 | 0.328 | 0.321 | 0.321 | 0.343 | 0.303 | 0.323 | 0.297 |
| AC-like 2 | 0.340 | 0.241 | 0.220 | 0.407 | 0.431 | 0.354 | 0.342 | 0.340 | 0.330 | 0.340 | 0.319 | 0.325 | 0.336 | 0.315 |
| Neu- like | 0.198 | 0.208 | 0.181 | 0.258 | 0.293 | 0.239 | 0.385 | 0.386 | 0.358 | 0.374 | 0.352 | 0.379 | 0.380 | 0.375 |
| Glycolysis_hi | 0.282 | 0.245 | 0.223 | 0.351 | 0.394 | 0.310 | 0.394 | 0.396 | 0.377 | 0.392 | 0.349 | 0.371 | 0.384 | 0.365 |

**Supplementary Figure 4.** Heatmap of cell type identification enrichment analysis for **A)** PA and **B)** GG subclusters. High values (red) indicate that many cells are shared between two clusters while low values (blue) indicate there is little overlap.

**Supplementary Figure 5**

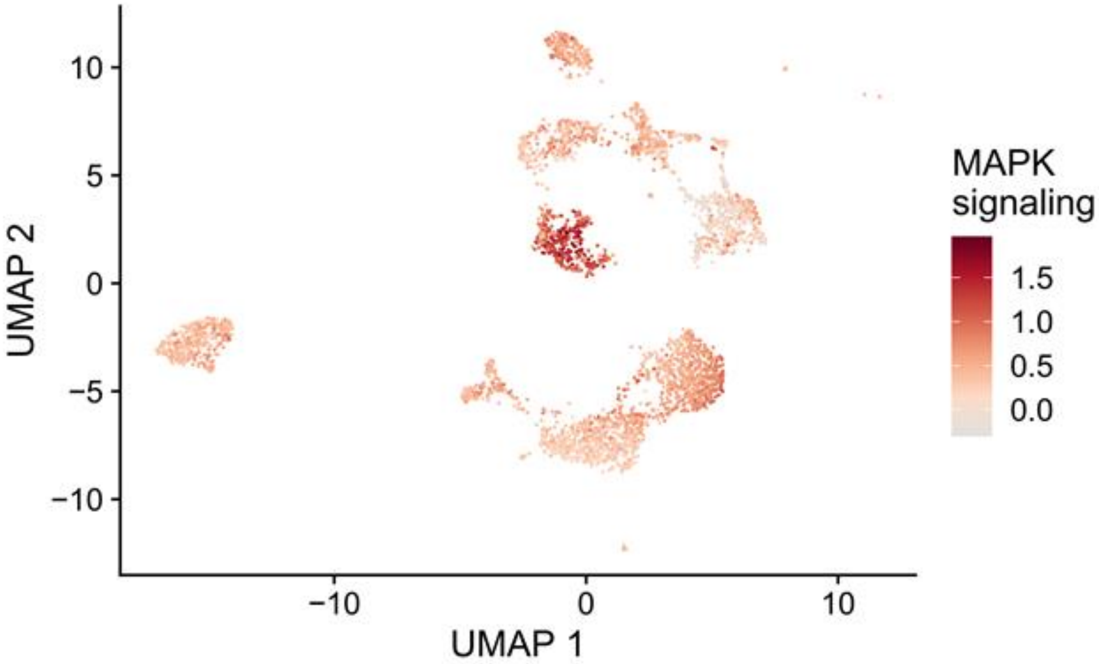


**Supplementary Figure 5.** UMAP plot demonstrating a subcluster of cells showing high MAPK activity in the PA dataset.

#### Supplementary Figure 6

A

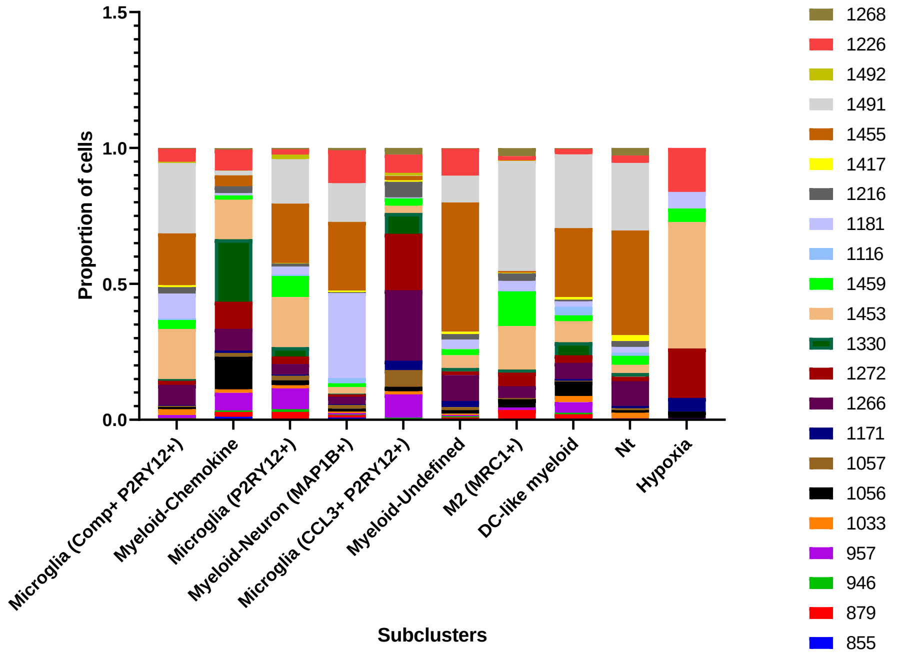


B

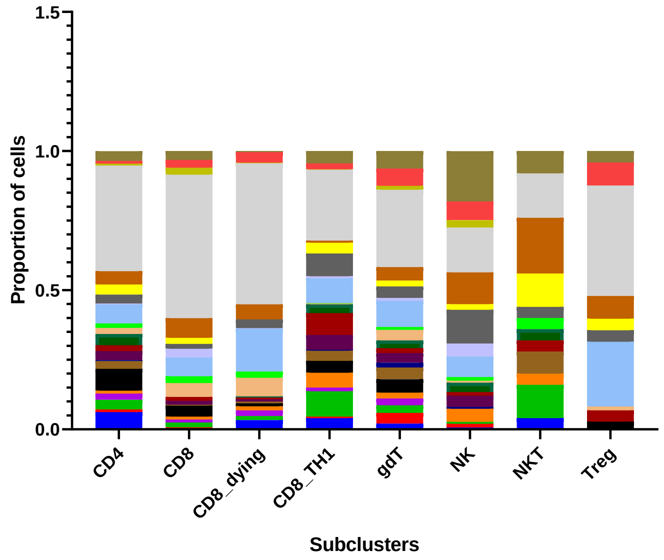


**Supplementary Figure 6.** Stacked barcharts show the contribution of each patient sample to various **A)** Myeloid and **B)** T cell subclusters. Cell counts for subcluster are normalized to 1.

### Supplementary Figure 7

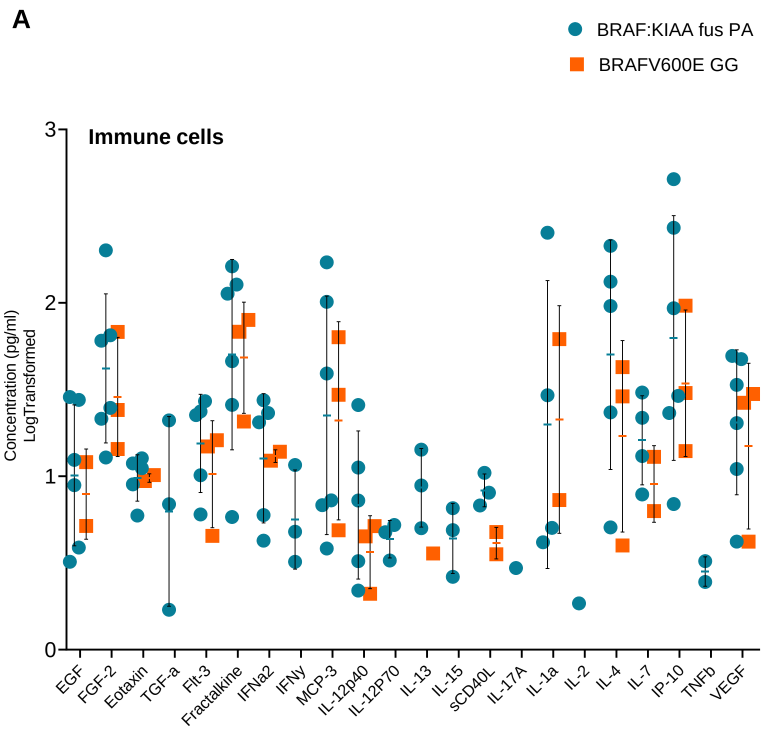


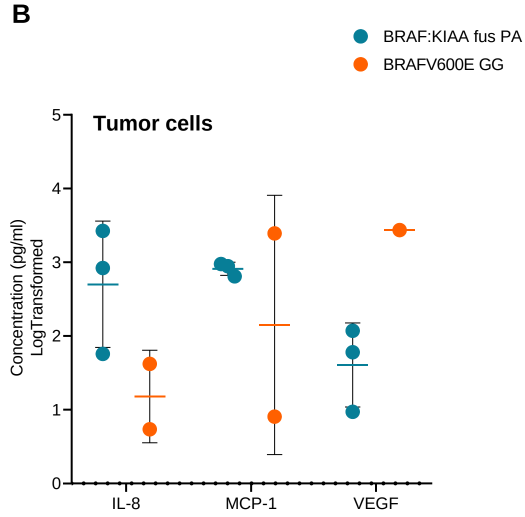


**Supplementary Figure 7.** Dot plots of detected cytokines/chemokines from bulk **A)** myeloid (all cytokines/chemokines) and **B)** tumor (cytokines/chemokines with concentrations greater than 1000pg/ml) cells conditioned media in PA and GG.

**Supplementary Figure 8.**

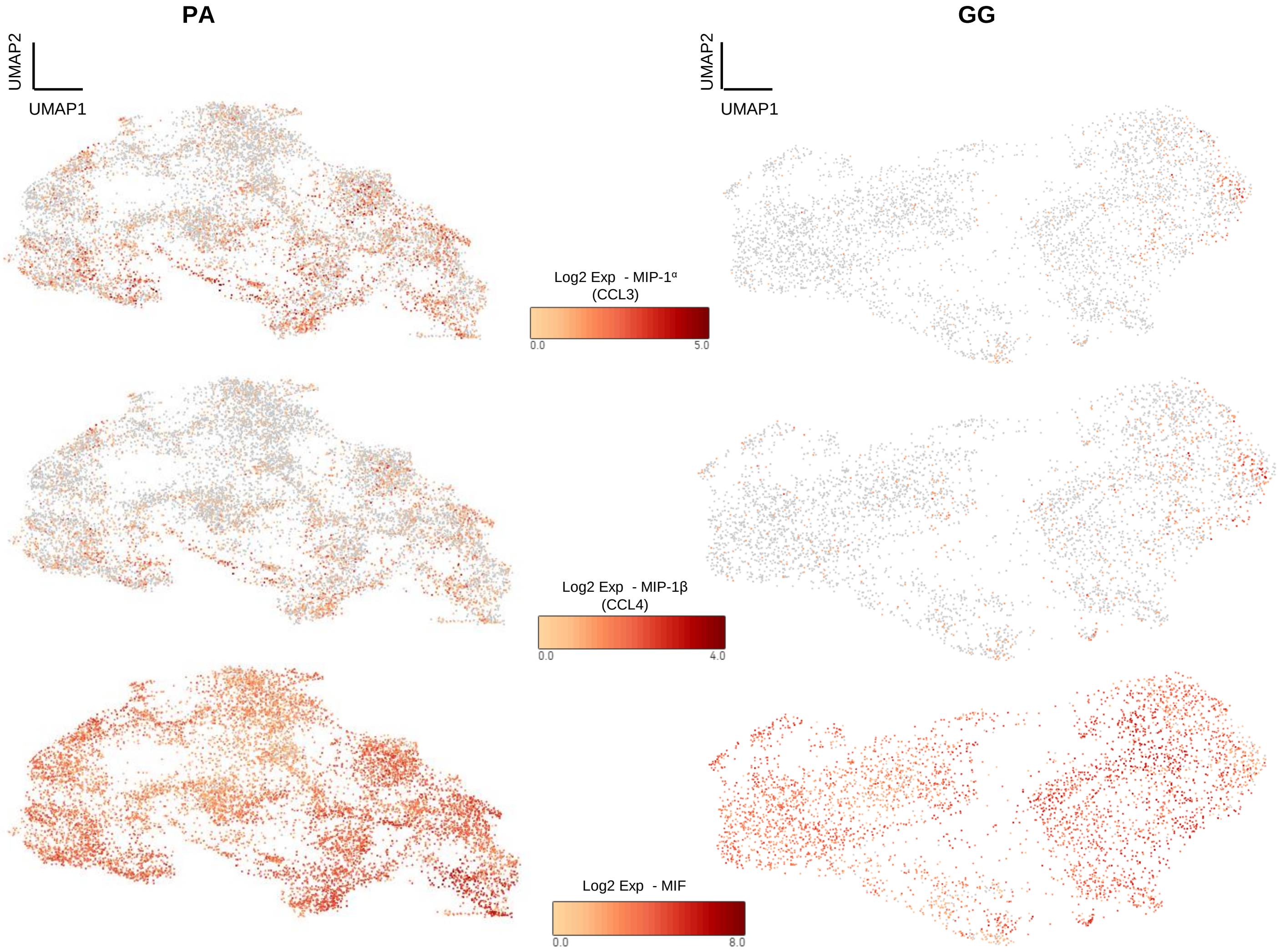


**Supplementary Figure 8.** UMAPs demonstrating spatial expression of MIP-1α (CCL3), MIP-1β (CCL4), and MIF in the PA and GG dataset.
